# Unilateral and Bilateral Subthalamic stimulation differently promote apathy: a translational approach

**DOI:** 10.1101/2020.06.11.147116

**Authors:** Yvan Vachez, Marie Bahout, Robin Magnard, Pierre-Maxime David, Carole Carcenac, Gabriel Robert, Marc Savasta, Sebastien Carnicella, Marc Vérin, Sabrina Boulet

**Affiliations:** Inserm, U1216, F-38000 Grenoble, France; Univ. Grenoble Alpes, Grenoble Institut des Neurosciences, GIN, F-38000 Grenoble, France; Behavior and Basal Ganglia research unit, University of Rennes 1-Rennes University Hospital, France; Neurology Department, Pontchaillou Hospital, Rennes University Hospital, France

**Author notes:** Equal contribution (Yvan Vachez, Marie Bahout). Co-senior author (Marc Vérin, Sabrina Boulet). **Correspondence:** Dr. Sabrina Boulet, INSERM U1216, Grenoble Institute of Neuroscience, Group “Pathophysiology of Motivation”, Grenoble Université - Site Santé La Tronche - BP 170, 38042 Grenoble, France., Pr. Marc Vérin, Behavior and Basal Ganglia research unit, University of Rennes 1-Rennes University Hospital, France.

## Abstract

Apathy, depression, and anxiety represent the main neuropsychiatric symptoms of Parkinson’s disease (PD). How subthalamic nucleus deep brain stimulation (STN-DBS) influences these symptoms, especially apathy, is a controversial topic. The present translational study investigates and compares the effect of bilateral or unilateral STN-DBS on this neuropsychiatric triad, combining a pre-clinical approach in rodents and a clinical follow-up of patients with PD. While depression and anxiety related behaviors remain unchanged, bilateral but not unilateral STN-DBS consistently induces apathy in patients and a reward seeking deficit in rodents. Together, these data substantiate the claim that STN-DBS may induce apathy by itself and suggest that bilateral but not the unilateral stimulation might be a critical factor.

## Introduction

Subthalamic nucleus (STN) deep brain stimulation (DBS) is a well-established alternative treatment for patients with Parkinson’s disease (PD) for whom classical pharmacological treatments have become ineffective(Benabid *et al*., 2009). According to the severity of the disease and the presence or lack of bilateral symptoms, STN-DBS is applied in an unilateral or bilateral fashion and permits physicians to efficiently alleviate motor disturbances(Samii *et al*., 2007; Benabid *et al*., 2009). Despite its manifest therapeutic benefits, STN-DBS effects on PD non-motor symptoms still represent a grey area (Ortega-Cubero *et al*., 2013; Fabbri *et al*., 2017). Numerous emotional, behavioral, and cognitive sideeffects are frequently reported (Castrioto *et al*., 2014). Apathy, depression, and anxiety represent the core neuropsychiatric symptoms of PD (Chaudhuri and Odin, 2010). Apathy is characterized by a loss of motivation or a reduction in goal-directed behaviors (Marin, 1996; Aarsland *et al*., 2009; Chaudhuri and Odin, 2010). It is certainly the symptom for which STN-DBS impact has been the most controversial (Czernecki *et al*., 2005; Drapier *et al*., 2006) and the debate is still ongoing (Zoon *et al*., 2021). While depression and anxiety do not seem consistently impacted by STN-DBS (Lhommee *et al*., 2012; Mansouri *et al*., 2018), apathy induction or aggravation following STN-DBS is frequently reported, reducing the quality of life improvement (Le Jeune *et al*., 2009; Castrioto *et al*., 2014; Martinez-Fernandez *et al*., 2016; Zoon *et al*., 2020). To explain this side-effect, one hypothesis is the resurgence of pre-existing apathy revealed by the classical reduction of dopatherapy following STN-DBS (Thobois *et al*., 2010; Chagraoui *et al*., 2018). The other hypothesis is the pro-apathetic effect of STN-DBS itself supported by clinical reports of apathy after bilateral STN-DBS which were not correlated with the reduction of the pharmacotherapy (Drapier *et al*., 2006; Drapier *et al*., 2008; Le Jeune *et al*., 2009; Robert *et al*., 2014; Zoon *et al*., 2019; Boon *et al*., 2021). We confirmed these clinical observations, showing that chronic and uninterrupted bilateral STN-DBS in naïve rats induces a reward seeking deficit, hypoactivity and exacerbate those symptoms already present in a model of PD neuropsychiatric symptoms (Vachez *et al*., 2020; Vachez and Creed, 2020).

Although we demonstrated the pro-apathetic effect of STN-DBS applied bilaterally, the unilateral STN-DBS effect remains to be investigated. Here, we provide both preclinical and clinical data in a translational investigation. We test whether unilateral STN-DBS induces a loss of motivation in rats, and we compare those results with a longitudinal follow-up of apathy incidence in unilaterally STN stimulated patients. We provide a comparison with the preclinical (Vachez *et al*., 2020) and clinical (Le Jeune *et al*., 2009) counterparts in studies we previously achieved using bilateral STN-DBS, conducted by the same centers and by the same clinical and scientific groups.

## Results and Discussion

Bilateral STN-DBS is applied in patients suffering from advanced stage PD with bilateral symptoms whereas unilateral STN-DBS is advocated for unilateral or very asymmetrical motor symptoms. Few clinical studies compared unilateral and bilateral STN-DBS, but those investigation consistently highlight that bilateral STN-DBS seems more effective In alleviating motor disturbances such as gait, tremor and oculomotor control (Huss *et al*., 2015; Lizarraga *et al*., 2016; Goelz *et al*., 2017). However, it is yet uncertain if the bilateralism of STN-DBS is critical for non-motor side effects, in particular apathy.

### Bilateral but not unilateral subthalamic stimulation induces a sustainable loss of motivation in rats

Because apathy particularly affects activity of daily living and simple tasks, motivation in rats was assessed with a sucrose fixed ratio 1 self-administration task. We apply STN-DBS with a portable microstimulator allowing stimulation 24 hours a day during several weeks. Bilateral STN-DBS induces an acute and persistent deficit in this task **(Figure 1A and B)**. However, unilateral STN-DBS acutely induces a strong decrease of reward seeking the first day, but this deficit does not last and is only transient and progressively disappears with the chronicity of STN-DBS **(Figure 1C and D)**. Furthermore, while the sucrose self-administration deficit induced by bilateral STN-DBS aligns with a hypoactive phenotype, highlighted by an open area task **(Figure 2A)**, unilateral STN-DBS does not induce behavioral hypoactivity, with both control and stimulated groups travelling the same distance during the task **(Figure 2B)**. However, left and right limbs fine motor skills during a stepping test right after the first session of sucrose self-administration are not altered either by bilateral **(Figure 2C)** or unilateral **(Figure 2D)** STN-DBS. Overall, only bilateral STN-DBS reduced motivated behaviors in rats. Stimulation parameters cannot account for this difference since we used similar current amplitude (Bilateral STN-DBS: 183 ± 13 μA; unilateral: 176 ± 11 μA).

**Figure 1:**
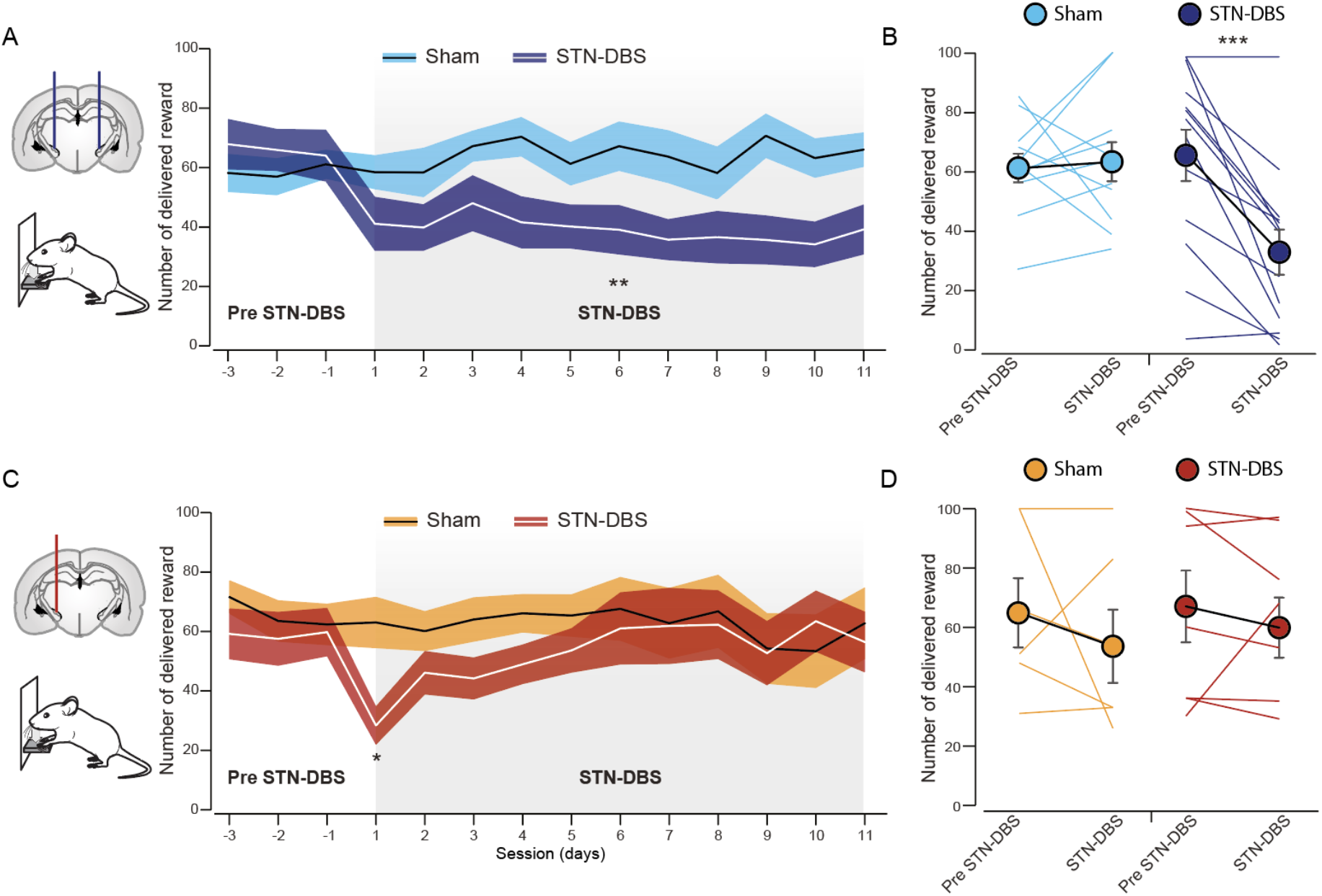
Unilateral STN-DBS does not reduce reward seeking. **(A)** and **(B)** Bilateral STN-DBS consistently decreased reward seeking. **(A)** Time course of the sucrose self-administration experiment. STN-DBS effect appears the first day of stimulation and this effect is stable and sustainable (RM ANOVA: Sessions effect: F_5.472,118.2_ = 0.7904, *P* = 0.5688; STN-DBS, F_1,22_ = 8.062, *P* = 0.0095). **(B)** Mean of the last 3 days (Paired t test, Sham: t_10_ = 0.2964, *P* = 0.7730; STN-DBS: t_12_ = 4.876, *P* = 0.0004). **(C)** and **(D)** Unilateral STN-DBS does not decrease reward seeking in a sustainable way. **(C)** Time course of the sucrose self-administration experiment. Unilateral STN-DBS only induce an acute decrease of the number of sucrose deliveries but this is not sustained throughout days (RM ANOVA: STN-DBS effect: F_1,27_ = 1.206, *P* = 0.2819; Sessions effect: F_10,179_ = 2.466 *P* = 0.0482; STN-DBS x Session interaction: F_10,179_ = 2.767, *P* = 0.0034). **(D)** Mean of the last 3 days. (Paired t test, Sham: t_5_ = 0.7815, *P* = 0.4699; STN-DBS: t_6_ = 0.02022, *P* = 0.9845). Bilateral: Sham: n = 13, STN-DBS: n = 13; Unilateral: Sham: n = 6, STN-DBS: n = 7 Data shown as means ± SEM. * *P* < 0.05; ** *P* < 0.01; *** *P* < 0.001.

**Figure 2:**
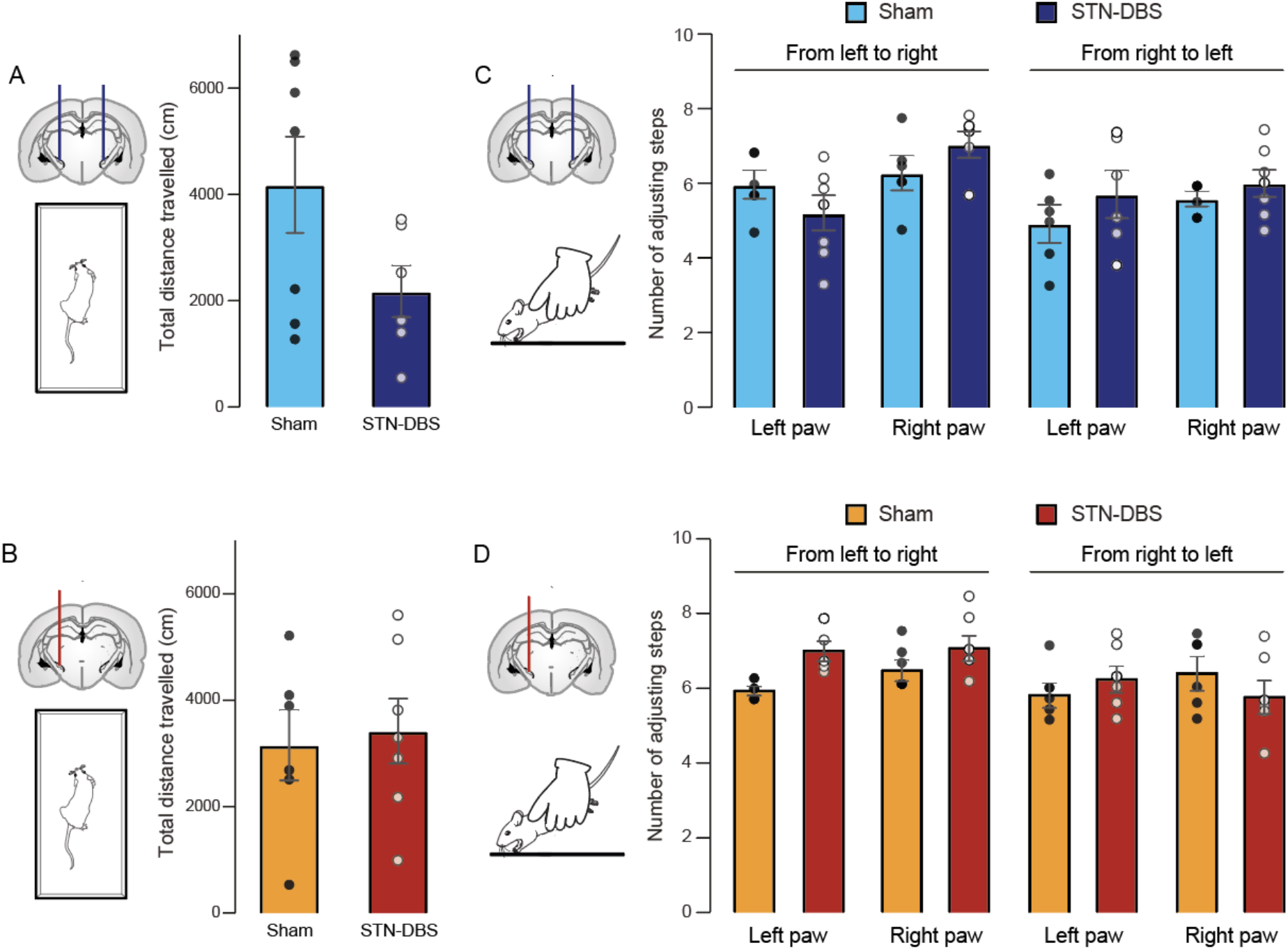
Unilateral STN-DBS does not alter basal locomotor activity or motor skills. **(A)** Bilateral STN-DBS tends to induce hypoactivity in rats during an open area test and decreases the total travelled distance (t test, t_11_ = 1.853, *P* = 0.0908). **(B)** Unilateral STN-DBS does not modify behavioral activity an open area test (t test, t_11_ = 0.2932, *P* = 0.7748) Neither **(C)** bilateral STN-DBS (RM 1 way ANOVA, STN-DBS effect: F_1,3_ = 0.6834, *P* = 0.4690; trials effect: F_1,1_ = 2.533, *P* = 0.3571) nor **(D)** unilateral STN-DBS (RM 1 way ANOVA, STN-DBS effect: F_1,3_ = 0.9981, *P* = 0.3914; trials effect: F_1,1_ = 0.9529, *P* = 0.5077; alter the fine motor skills of front limbs as demonstrated by adjusting steps in the course of the stepping test. Bilateral: Sham: n = 7, STN-DBS: n = 6; Unilateral: Sham: n = 6, STN-DBS: n = 7 Data shown as means ± SEM.

### Subthalamic stimulation does not promote anhedonic or anxiety related behaviors In rats

Alongside apathy, depression, and anxiety are the main neuropsychiatric symptoms present in PD patients (Chaudhuri *et al*., 2006). Anhedonia is a core feature of depression and is defined as the inability to experience pleasure (Assogna *et al*., 2011; Rizvi *et al*., 2016). We evaluated the impact of STN-DBS on this behavior with a two-bottle choice procedure. Neither bilateral **(Figure 3A)** nor unilateral **(Figure 3B)** modify the preference of sucrose over water, meaning that hedonic behaviors are unchanged by STN-DBS. Those results seem contradictory since bilateral STN-DBS in rats has been shown to induce depressive like behavior (Temel *et al*., 2007; Creed *et al*., 2013). However, the modality of the stimulation (polarity, chronicity, etc), along with the difference between the tests can explain those discrepancies (as discussed in Vachez and Creed, 2020).

**Figure 3:**
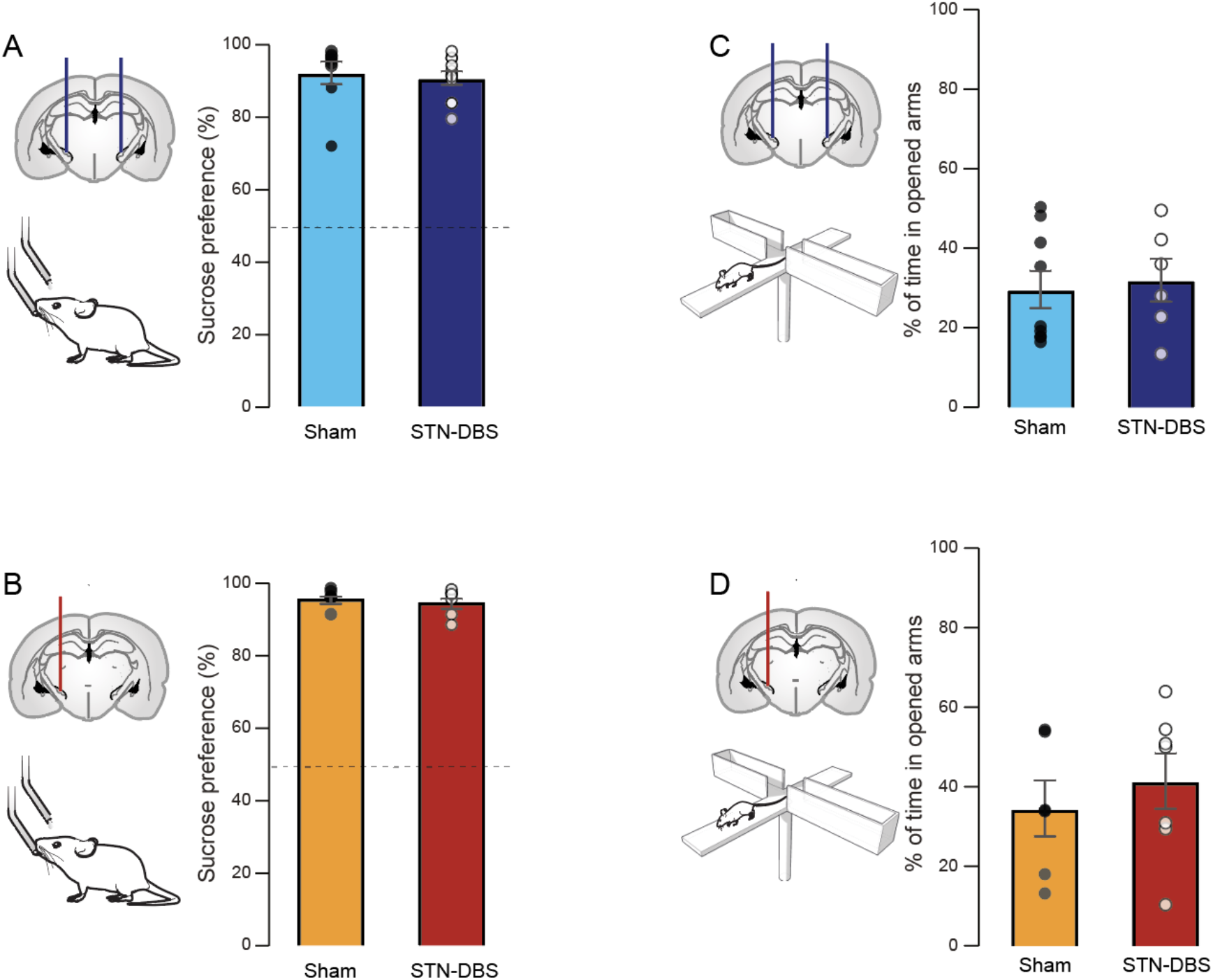
Bilateral or unilateral STN-DBS do not promote anhedonic or anxiety related behavior. **(A)** Bilateral and **(B)** unilateral STN-DBS do not alter preference for sucrose over water in a two-bottle choice test (Bilateral STN-DBS: t_17_ = 0.5485, *P* = 0.5943; Unilateral STN-DBS: t_11_ = 0.5485, *P* = 0.5943). **(C)** Bilateral and **(D)** unilateral STN-DBS do not modify the time spent in the open arms of an elevated plus maze (Bilateral STN-DBS: t_13_ = 0.3195, *P* = 0.7545; Unilateral STN-DBS: t_11_ = 0.5084, *P* = 0.5084). Bilateral: Sham: n = 8, STN-DBS: n = 11; Unilateral: Sham: n = 6, STN-DBS: n = 7 Data shown as means ± SEM.

Next, while the absence of effect of bilateral STN-DBS on anxiety related behavior in an elevated plus maze has previously been demonstrated (Creed *et al*., 2013), we replicate that result **(Figure 3C)** and show in addition that unilateral STN-DBS exerts the same null effect on the time spent in the open arms **(Figure 3D)**.

Altogether, these data show that, in the exact same condition, only bilateral STN-DBS induces a motivational deficit, reminiscent of apathy in stimulated PD patients whereas anhedonic or anxiety behaviors are not altered by STN-DBS, either unilateral or bilateral.

### Bilateral but not unilateral subthalamic stimulation promotes apathy in Parkinson’s disease patients

Few clinical studies compared unilateral and bilateral STN-DBS in same patients, but it is admitted that bilateral STN-DBS induces greater therapeutic effect (Samii *et al*., 2007). However, bilateral STN-DBS seems also to induce more deleterious side effects, aggravating cognitive processes, loss of verbal fluidity and weight gain in a greater proportion than unilateral stimulation (Lee *et al*., 2011; Sjoberg *et al*., 2012; Goelz *et al*., 2017). We previously made the unique description of post-STN-DBS apathy in bilaterally stimulated patients, that was not correlated to dopaminergic reduction (Drapier *et al*., 2006; Drapier *et al*., 2008; Le Jeune *et al*., 2009; Robert *et al*., 2014). Here, we are providing the follow-up of patients who are treated in our center in the exact same conditions but with unilateral STN-DBS. Contrary to bilateral stimulation, unilateral STN-DBS does not increase the mean apathy evaluation scale score or the number of apathetic patients **(Figure 4A)**. Furthermore, despite a deleterious tendency of bilateral STN-DBS to increase depression (MADRS score) and anxiety (AMDP-AT score), both symptoms are not affected by unilateral STN-DBS **(Figure 4B and 4C)**.

**Figure 4:**
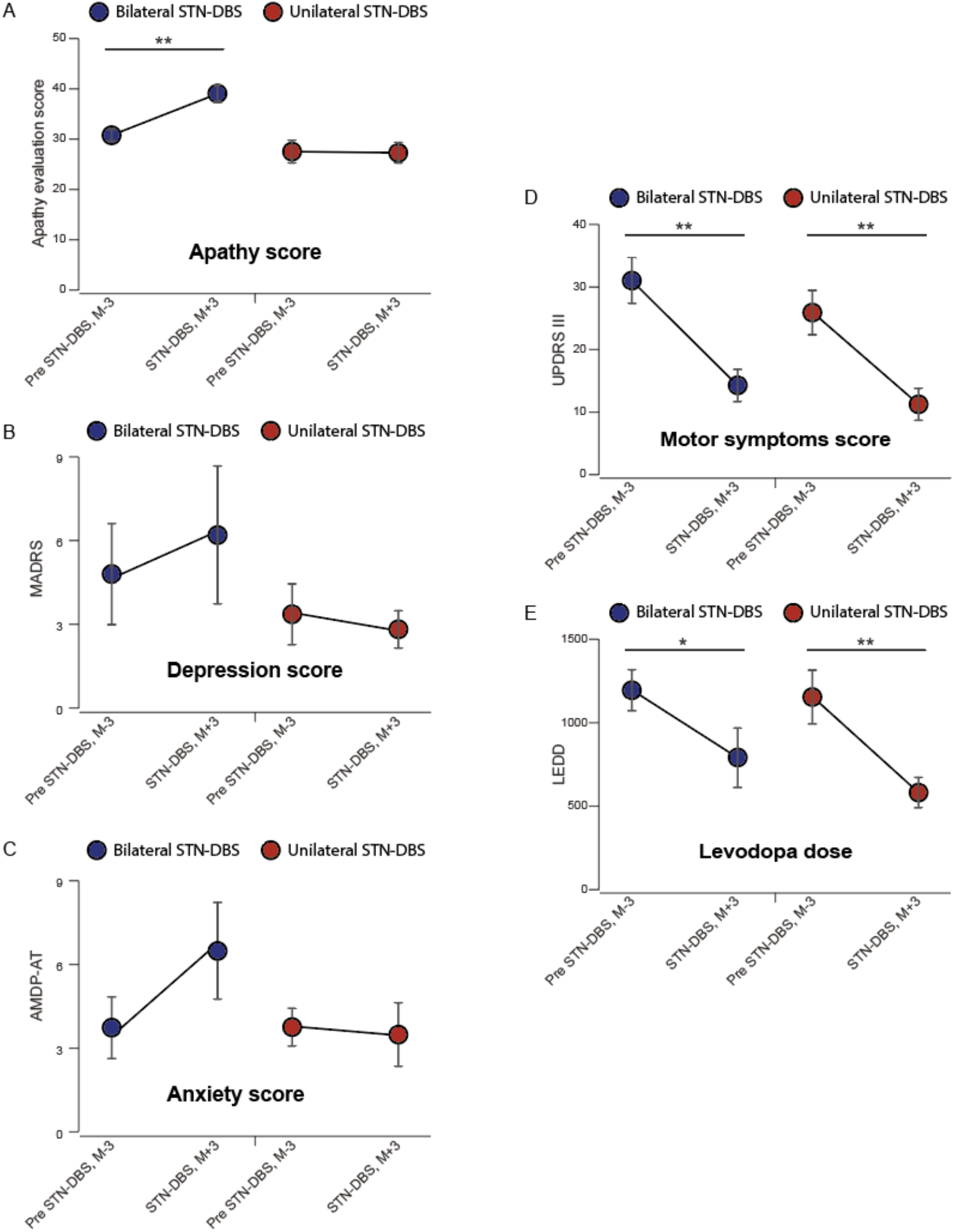
Bilateral but not unilateral STN-DBS promotes apathy in Parkinson’s disease patients. **(A)** Bilateral STN-DBS promotes apathy, highlighted by a significant increase of the Apathy Evaluation Scale (AES) score (Wilcoxon signed rank test, *P* = 0.002) but not unilateral STN-DBS (Wilcoxon signed rank test, *P* = 0.82). **(B)** Bilateral or unilateral STN-DBS do not alter the Montgomery-Åsberg Depression Rating Scale (MADRS) score (Wilcoxon signed rank test, bilateral STN-DBS: *P* = 0.5, unilateral STN-DBS: *P* = 0.64). **(C)** Bilateral or unilateral STN-DBS do not significantly alter the Anxiety Scale of Association for Methodology and Documentation in Psychiatry (AMDP-AT) score (Wilcoxon signed rank test, bilateral STN-DBS: *P* = 0.2, unilateral STN-DBS: *P* = 0.8). **(D)** Bilateral and unilateral STN-DBS permit a similar Unified Parkinson’s Disease Rating Scale III (UPDRS III) score improvement (Wilcoxon signed rank test, bilateral STN-DBS: *P* = 0.007, unilateral STN-DBS: *P* = 0.0011)**. (E)** The Levodopa-Equivalent Daily Doses (LEDD) and their reduction were similar between patients undergoing bilateral or unilateral STN-DBS (Wilcoxon signed rank test, bilateral STN-DBS: *P* = 0.02, unilateral STN-DBS: *P* = 0.0011). Bilateral: n = 12; Unilateral: n = 11. Data shown as means ± SEM. * *P* < 0.05; ** *P* < 0.01.

This absence of post unilateral STN-DBS apathy cannot be attributed to an ineffective stimulation or electrodes misleading, since all patients present a significant postoperative motor improvement in OFF and ON-dopa conditions, shown by UPDRS II to IV, the Hoehn and Yahr and the Schwab and England scores **(Figure 4D and Supplemental Table 1)**. Furthermore, the pre-STN-DBS Levodopa-Equivalent Daily Dose (LEDD) (1162.0 ± 533.5 mg) and its reduction following the surgery (589.0 ± 302 mg) match the dose prescribed to the cohort of patients who developed apathy following bilateral STN-DBS (from 1200.0 ± 426.5 mg to 796.66 ± 620.0 mg) **(Figure 4E)** (Le Jeune *et al*., 2009). Finally, even if the number of patients (n=11) is relatively low, it is unlikely that this prevents us from detecting an apathetic effect of unilateral STN-DBS, since we were able to statistically highlight apathy increases following bilateral STN-DBS in two previous studies with small cohorts from the same clinical groups (Drapier *et al*., 2006; Le Jeune *et al*., 2009).

A main limitation of this clinical follow-up is the absence of random assignment; such a study would be unethical because patients with bilateral symptoms would not being given the best possible chances with unilateral STN-DBS and conversely patients with unilateral or very asymmetrical symptoms would be exposed to an unjustified risk with a bilateral STN-DBS.

Overall, in addition to provide a significant clinical improvement, unilateral STN-DBS on patients seems to be less pathogenic regarding post-operative apathy compared to bilateral STN-DBS. However, our preclinical data suggest that unilateral STN-DBS in patients could have acutely promote apathy, but that effect progressively disappeared and was absent at 3 months post STN-DBS, when apathy was assessed. This suggests the implementation of progressive contralateral compensatory mechanism which may involve inter hemispheric connection between both STN (Mark *et al*., 2005; Hohlefeld *et al*., 2014), in addition to contralateral connectivity with several input or output structures (Lanciego *et al*., 2012). To this extent, unilateral lesion of the substantia nigra pars compacta induces in rats a characteristic and dramatic increase of the firing rate of the ipsilateral STN, but conversely decreases the activity of the contralateral STN (Périer *et al*., 2000). Furthermore, while it is commonly described that STN-DBS locally inhibits the activity of STN, unilateral STN-DBS in patients is associated with increased activity of the contralateral STN (Novak *et al*., 2009; Walker *et al*., 2011).

Although bilateral STN-DBS has quickly become a standard way to apply DBS, this should not be an exclusive approach for all PD patient candidates for stimulation. Efficacy and safety of unilateral STN DBS has been established in PD for patients with unilateral (Kumar et al., 1999) or bilateral asymmetric motor symptoms (Walker et al., 2009), with even significant effect on the ipsilateral hemi body. In practice, a unilateral surgery could be a good option for young patients with prominent asymmetric symptoms in order to protect them from apathy. Later, if necessary, a contralateral surgery could be a reasonable option.

To summarize, the present translational work refines the importance of STN-DBS modality (unilateral versus bilateral) in the induction of apathy. Hence, it legitimizes the reconsideration of the rationale for unilateral versus bilateral STN-DBS use in PD patients regarding unilateral STN-DBS efficiency together with the lower incidence of non-motor sideeffect, including apathy.

## Methods

### Experimental study

Protocols used for the experimental part complied with the European Union 2010 Animal Welfare Act and the French directive 2010/63.

#### Animals

Experiments were performed on 80 adult male Sprague-Dawley rats (Janvier, Le Genest-Saint-Isle, France) weighing approximately 350g (8 weeks old) at the time of surgery. Animals were individually housed under standard laboratory conditions (12 h light/dark cycle, with lights on at 7 a.m) with food and water available *ad libitum*. Protocols used complied with the European Union 2010 Animal Welfare Act and the French directive 2010/63.

#### Unilateral implantation of electrodes in the subthalamic nucleus

Rats were unilaterally implanted in left or right STN with monopolar electrodes consisting of platinum-iridium wire insulated with Teflon with a 400 μm exposed end (wire diameter, 110 μm insulated, 76 μm bare, PT-IR Teflon, Phymep, Paris, France). Stereotaxic coordinates (relative to bregma, according to the stereotaxic atlas of Paxinos and Watson(Paxinos G, 1998)) were: AP, −3.8 mm; L, ±2.4 mm, and DV, −7. 8 mm; with the incisor bar at +3.2 mm below the interaural plane. The exposed end of the electrode, located in the STN, corresponded to the negative stimulation pole. A screw (Phymep Paris, France, 0-80×1/16) fixed on the skull was used as the positive pole. Electrodes were soldered to corresponding contacts of the microstimulator support (ISENUSTIMV7, ISEN, Toulon, France) which was permanently fixed to the rat skull with dental cement (Superbond, Phymep Paris, France). After recovery from anesthesia, animals were returned to the facility for 3 days, to allow recovery before the beginning of the behavioral experiments.

#### Subthalamic nucleus deep brain stimulation

Chronic long-lasting unilateral STN-DBS was performed using an electrical portable microstimulator system already validated in freely moving rats(Forni *et al*., 2012; Vachez *et al*., 2020). This system has the advantages of leaving the animals free to move during behavioral tasks, of being removable and of allowing rapid and easy activation (ON/OFF), modulation of DBS parameters or battery change without strain of rats or anesthesia.

The interface between electrodes and the microstimulator is a support with the top side designed as a platform to receive a microstimulator’s plug in. The microstimulator is made up with classical structures allowing to regulate the frequency and the duty cycle. A varying resistance provides a current peak with variable current pulse amplitudes. The power supply of the microstimulator consists of two 3 V flat lithium watch batteries (CR1220) connected in series. The generator micro-circuit is coated with a polyurethane resin (UR5041 Electrolub).

The frequency and pulse width used, 130 Hz and 60 μs respectively, were similar to those applied in humans. For each animal, intensity was gradually increased until dyskinetic movement of the contralateral forelimb appeared, as described by Boulet et al.(Boulet *et al*., 2006) and then adjusted just below this pro-dyskinetic threshold. This value was conserved throughout the study. The stimulation intensity was 176 +/- 11 μA on average and ranged between 100 and 225 μA. These parameters were determined the day before the beginning of stimulation. STN-DBS was uninterrupted until rat euthanasia.

#### Behavioral procedures

Rats were not food or water deprived during the experimental procedures.

##### Sucrose self-administration

Rats were trained to work for a 2.5% sucrose reinforcer, chosen for its relatively moderate rewarding outcome, in a self-administration task in operant chambers (Med Associates, St Albans, VT, USA) under a fixed ratio 1 reinforcement schedule. Rats were given the choice between two levers: an active and reinforced one, delivering 0.2 ml sucrose solution in a receptacle when pressed, and an inactive, non-reinforced lever, producing nothing(Carnicella *et al*., 2014; Drui *et al*., 2014; Favier *et al*., 2014; Favier *et al*., 2017).

Conditions were counter-balanced among the different test. Trainings occurred before and after electrodes implantation. Once performances were stable during the second training, (less than 20% performance variation over three consecutive sessions), STN-DBS was turned ON.

##### Two bottle choice

In their home cage, rats were given 24h concurrent access to two graduated 250 ml plastic bottles (Techniplast, Lyon, France), for 3 days. One bottle contained tap water, whereas the other contained 2.5% sucrose (Sigma) in tap water. Rats and bottles were weighed daily, with the position of the bottles (left or right) alternated to control for side preference. The first day was used as an acclimation period. The volumes of sucrose solution and water consumed on the second and third days were averaged to determine preference for sucrose over water (sucrose intake/total intake, expressed as a percentage).

##### Stepping test

Animals were moved sideways along a smooth-surfaced table over 90 cm and the number of forelimb adjusting steps measured(Olsson *et al*., 1995; Drui *et al*., 2014). The test was carried out three times for each paw.

##### Open area

Rats were placed in a dimly lit white Perspex open arena (50 x 25 x 40cm) and horizontal distances traveled were recorded with a video-tracking system to assess locomotor and basal activity (Viewpoint S.A., Champagne au Mont d’Or, France), over a 1h period.

##### Elevated plus maze

The elevated plus maze consists of two opposing open arms and two opposing arms enclosed by 40 cm high walls, and was placed in a dimly lit room. Each arm was 50 cm long and 10cm wide and made of black Perspext, suspended 55cm above the floor. The rats were placed in the center of the maze and their behavior was recorded for 5 min with a videotracking system. Total time spent in the open and closed arms were quantified by the videotracking system.

#### Histological analysis

Briefly, rats were sacrificed under chloral hydrate anesthesia at the end of the behavioral experiments, intracardially perfused with NaCl (0,9%) and their brains frozen in cooled isopentane (−40°C) and stored at −30°C. Serial coronal sections (14-μm thick) of subthalamic nucleus were cut with a cryostat (Microm HM 500, Microm, Francheville, France) and collected on slides.

Sections were stained with Cresyl violet and analyzed under a light microscope (Nikon, Eclipse 80i, TRIBVN, Châtillon, France) coupled to the ICS FrameWork computerized image analysis system (TRIBVN, 2.9.2 version, Châtillon, France) in order to check the positions of electrodes. Animals with incorrect electrode locations were excluded from the study.

### Clinical study

Oral and written informed consent was obtained for all subjects of the clinical study, which was consistent with ethical guidelines of Helsinki’s Declaration.

#### Patients

The group of subjects used in this study was composed of 11 adult patients who suffered of unilateral symptoms of Parkinson’s disease according to clinical criteria of UK Parkinson’s Disease Society Brain Bank for Idiopathic Parkinson’s Disease(Hughes *et al*., 1992), refractory to medical treatment, at the stage of severe ON/OFF fluctuations and dyskinesias. Nine men and two women underwent a unilateral STN DBS in Neurosurgery Department at Rennes University Hospital Center (France), between September 2006 and March 2016. All patients fulfilled selective criteria for this surgery, with unilateral or very asymmetric bilateral motor symptoms, no neuropsychological disorder and no brain atrophy or vascular lesions on preoperative MRI. Exclusion criteria for STN DBS were axial motor signs such as gait disorders or postural instability, psychiatric trouble, depression on the Montgomery-Åsberg Depression Rating Scale(Montgomery and Asberg, 1979), cognitive impairment on Mattis Dementia Rating Scale, executive functions trouble, which were not found for all patients(Welter *et al*., 2002). All psychiatric and neuropsychological assessment at inclusion were realized in on-dopa condition, accordingly to our previous studies.

Six patients underwent a left STN DBS while five patients underwent a right STN DBS. Single electrode was implanted in contralateral STN to affected hemibody; patients with right motor symptoms were implanted in left STN and patients with left symptoms in the right side. At time of surgery, mean age (± SD) of patients was 49.2 (± 9.0) years. The mean disease duration (± SD) was 7.3 (± 1.4) years. One patient was left-handed and ten were right-handed, accordingly to the Edinburgh Handedness Inventory criteria(Oldfield, 1971). Oral and written informed consent was obtained for all subjects of this study, which was consistent with ethical guidelines of Helsinki’s Declaration.

#### Neurosurgery and deep brain stimulation

Quadripolar electrodes of DBS were stereotactically implanted in brain patients (model 3389, Medtronic, Minneapolis, MN, USA). After 3D MRI and 3D computer tomography (CT), exact location in STN were determined for precision surgery, with a similar methodology described in previous study(Benabid *et al*., 2000). Intervention was realized on local anesthesia, with Leksell frame, associated with motor and vision testing to ensure the correct position of electrode and the absence of side effect with the lowest voltage. An implantation of pulse generator was also realized in left infraclavicular subcutaneous position. Stimulation was switched on in the few days following. A second 3D CT was realized three days after DBS surgery to confirm the correct electrode location in the STN. Monopolar stimulation with one contact of electrode was used for all patients. For the selected contact, the mean coordinates for patients with left STN were 12.1 ± 1.3 mm lateral to the anterior commissure- posterior commissure line, 0.9 ± 1.7 mm below the antero-posterior commissure- and 1.3 ± 1.4 mm posterior to the middle anteror-posterior commissure. The mean coordinates for patients with right STN were 11.5 ± 1.2 mm lateral to the antero-posterior commissure line, 0.9 ± 1.5 mm below the antero - posterior commissure and 1.6 ± 1.2 mm posterior to the middle anteroposterior commissure. The mean impulse duration (± SD) was 62.7 μs (± 9.0). The mean frequency (± SD) was 145 Hz (± 25.8). The mean voltage (± SD) was 2.6 V (± 0.6).

#### Clinical evaluation

The eleven patients were evaluated before surgery of DBS (3 months before) and after surgery (3 months after) using psychiatric, neuropsychological and motor scales. Motor evaluations were realized in OFF and ON-dopa conditions, while neuropsychological and psychiatric evaluations were realized in on-dopa and on stimulation conditions. Each patient underwent a regular follow-up for optimization of DBS-parameters by a neurologist specialized for Parkinson’s disease.

##### Psychiatric assessment

Patients were also subjected to a psychiatric evaluation with the clinician version of Apathy Evaluation Scale (C-AES)(Marin *et al*., 1991), the Montgomery-Åsberg Depression Rating Scale (MADRS) and the Anxiety Scale of Association for Methodology and Documentation in Psychiatry (AMDP-AT)(Bobon, 1985; Bobon *et al*., 1985) during on-drug condition, 3 months before and after neurosurgery.

The AES contains 18 questions with scores ranging from 18 to 72, and the highest score reflecting severe apathy. It is recognized as the most psychometrically robust apathy scale across any population and was given “suggested scale” status for PD(Leentjens *et al*., 2008). To define clinical apathy a cut-off score of 42 or above is chosen.

The MADRS scale was chosen because of the predominance of the psychic items rather than the somatic items which permits to limit the interferences with parkinsonian’s symptoms. Patients were assessed by trained psychiatrists sensitized to psychiatric disorders in PD.

The AMPDT AT was specifically chosen because of the little power of the somatic items.

##### Motor evaluation

All patients were evaluated according to the guidelines of the Core Assessment Program for Intracerebral Transplantation (CAPIT)(Langston *et al*., 1992). They were assessed before and after surgery by motor scores with the Unified Parkinson’s Disease Rating Scale (UPDRS) Part I to IV(Fahn, 1987), the Hoehn and Yahr scale(Hoehn and Yahr, 1967) and the Schwab and England scale(Schwab and England, 1969). These scores were evaluated in OFF-dopa (at least 12 hours without any dopa medication) and ON-dopa conditions with a dopamine challenge (with a dose equal to 50mg of levodopa added to morning dose). The total Levodopa-Equivalent Daily Dose (LEDD) of each patient was calculated before surgery and at the post-operative evaluation, accordingly to the basis method (Deuschl *et al*., 2006).

### Statistic

Statistics were performed in Graphpad prism version 8. A *P* value of 0.05 was considered significant. Experimental study: data are shown as means ± SEM. Student’s t-test was used for 2 groups analysis; otherwise, repeated measure or 2-way ANOVA were performed. Clinical study: data are shown as means ± SD. The Wilcoxon signed rank sum test was used to realize intra-group analyses.

---

**Supplemental Table 1.**
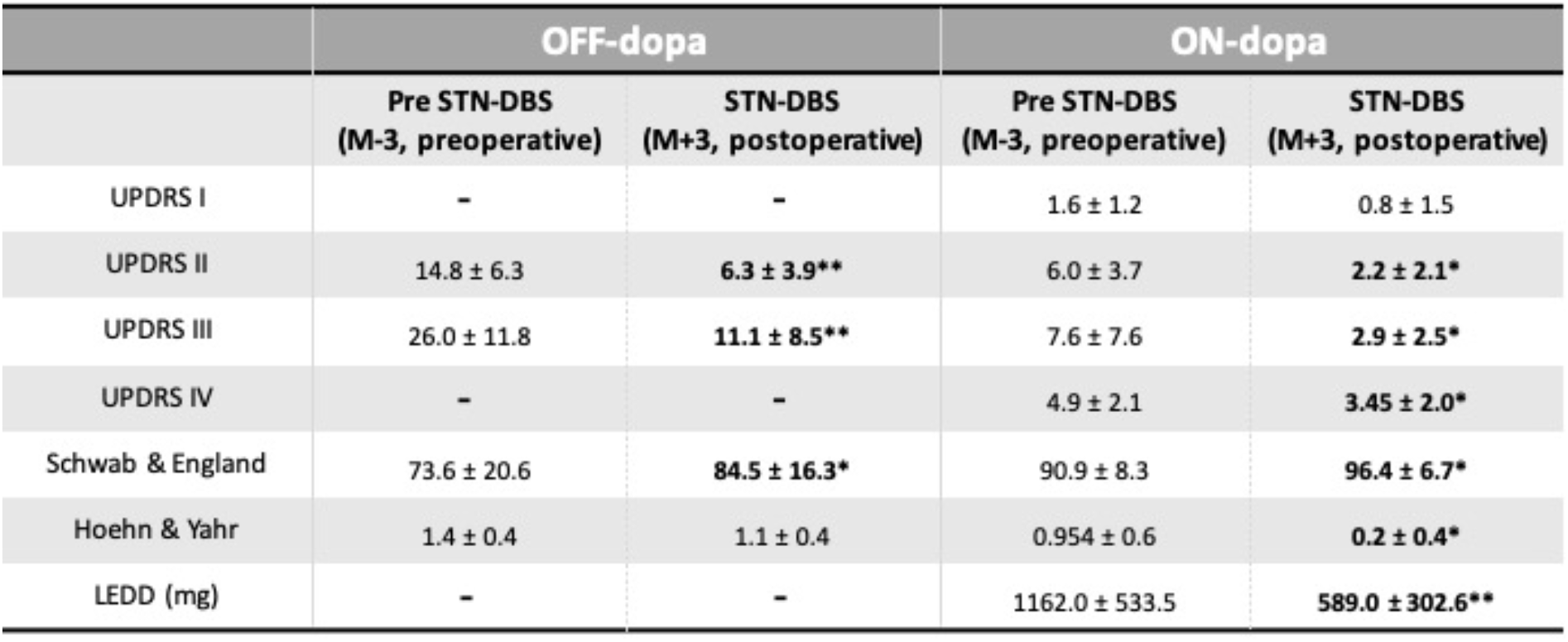
Clinical and motor scores (mean ± SD) of Parkinson’s disease patients 3 months before (M-3) and after (M+3) unilateral subthalamic nucleus stimulation. UPDRS: United PD Rating Scale; LEDD: Levodopa Equivalent Daily Dose.

## Notes

### Competing Interest Statement

The authors have declared no competing interest.

## References

Aarsland D, Bronnick K, Alves G, Tysnes OB, Pedersen KF, Ehrt U, et al. The spectrum of neuropsychiatric symptoms in patients with early untreated Parkinson’s disease. Journal of neurology, neurosurgery, and psychiatry 2009; 80(8): 928–30.

Assogna F, Cravello L, Caltagirone C, Spalletta G. Anhedonia in Parkinson’s disease: a systematic review of the literature. Movement disorders : official journal of the Movement Disorder Society 2011; 26(10): 1825–34.

Benabid AL, Chabardes S, Mitrofanis J, Pollak P. Deep brain stimulation of the subthalamic nucleus for the treatment of Parkinson’s disease. Lancet Neurol 2009; 8(1): 67–81.

Benabid AL, Krack PP, Benazzouz A, Limousin P, Koudsie A, Pollak P. Deep brain stimulation of the subthalamic nucleus for Parkinson’s disease: methodologic aspects and clinical criteria. Neurology 2000; 55(12 Suppl 6): S40–4.

Bobon D. [Contribution of the AMDP scale to quantitative psychopathology]. Acta psychiatrica Belgica 1985; 85(1): 5–249.

Bobon D, von Frenckell R, Troisfontaines B, Mormont C, Pellet J. [Preliminary construction and validation of an anxiety scale derived from the French version of the AMDP, the AMDP- AT]. L’Encephale 1985; 11(3): 107–11.

Boon LI, Potters WV, Zoon TJC, van den Heuvel OA, Prent N, de Bie RMA, et al. Structural and functional correlates of subthalamic deep brain stimulation-induced apathy in Parkinson’s disease. Brain Stimul 2021; 14(1): 192–201.

Boulet S, Lacombe E, Carcenac C, Feuerstein C, Sgambato-Faure V, Poupard A, et al. Subthalamic stimulation-induced forelimb dyskinesias are linked to an increase in glutamate levels in the substantia nigra pars reticulata. The Journal of neuroscience : the official journal of the Society for Neuroscience 2006; 26(42): 10768–76.

Carnicella S, Drui G, Boulet S, Carcenac C, Favier M, Duran T, et al. Implication of dopamine D3 receptor activation in the reversion of Parkinson’s disease-related motivational deficits. Transl Psychiatry 2014; 4: e401.

Castrioto A, Lhommee E, Moro E, Krack P. Mood and behavioural effects of subthalamic stimulation in Parkinson’s disease. Lancet Neurol 2014; 13(3): 287–305.

Chagraoui A, Boukhzar L, Thibaut F, Anouar Y, Maltete D. The pathophysiological mechanisms of motivational deficits in Parkinson’s disease. Prog Neuropsychopharmacol Biol Psychiatry 2018; 81: 138–52.

Chaudhuri KR, Healy DG, Schapira AH. Non-motor symptoms of Parkinson’s disease: diagnosis and management. The Lancet Neurology 2006; 5(3): 235–45.

Chaudhuri KR, Odin P. The challenge of non-motor symptoms in Parkinson’s disease. Progress in brain research 2010; 184: 325–41.

Creed MC, Hamani C, Nobrega JN. Effects of repeated deep brain stimulation on depressive- and anxiety-like behavior in rats: comparing entopeduncular and subthalamic nuclei. Brain Stimul 2013; 6(4): 506–14.

Czernecki V, Pillon B, Houeto JL, Welter ML, Mesnage V, Agid Y, et al. Does bilateral stimulation of the subthalamic nucleus aggravate apathy in Parkinson’s disease? Journal of neurology, neurosurgery, and psychiatry 2005; 76(6): 775–9.

Deuschl G, Schade-Brittinger C, Krack P, Volkmann J, Schafer H, Botzel K, et al. A randomized trial of deep-brain stimulation for Parkinson’s disease. The New England journal of medicine 2006; 355(9): 896–908.

Drapier D, Drapier S, Sauleau P, Haegelen C, Raoul S, Biseul I, et al. Does subthalamic nucleus stimulation induce apathy in Parkinson’s disease? Journal of neurology 2006; 253(8): 1083–91.

Drapier D, Peron J, Leray E, Sauleau P, Biseul I, Drapier S, et al. Emotion recognition impairment and apathy after subthalamic nucleus stimulation in Parkinson’s disease have separate neural substrates. Neuropsychologia 2008; 46(11): 2796–801.

Drui G, Carnicella S, Carcenac C, Favier M, Bertrand A, Boulet S, et al. Loss of dopaminergic nigrostriatal neurons accounts for the motivational and affective deficits in Parkinson’s disease. Mol Psychiatry 2014; 19(3): 358–67.

Fabbri M, Coelho M, Guedes LC, Rosa MM, Abreu D, Goncalves N, et al. Acute response of non-motor symptoms to subthalamic deep brain stimulation in Parkinson’s disease. Parkinsonism & related disorders 2017; 41: 113–7.

Fahn S. Drug treatment of hyperkinetic movement disorders. Seminars in neurology 1987; 7(2): 192–208.

Favier M, Carcenac C, Drui G, Vachez Y, Boulet S, Savasta M, et al. Implication of dorsostriatal D3 receptors in motivational processes: a potential target for neuropsychiatric symptoms in Parkinson’s disease. Sci Rep 2017; 7: 41589.

Favier M, Duran T, Carcenac C, Drui G, Savasta M, Carnicella S. Pramipexole reverses Parkinson’s disease-related motivational deficits in rats. Movement disorders : official journal of the Movement Disorder Society 2014; 29(7): 912–20.

Forni C, Mainard O, Melon C, Goguenheim D, Kerkerian-Le Goff L, Salin P. Portable microstimulator for chronic deep brain stimulation in freely moving rats. Journal of neuroscience methods 2012; 209(1): 50–7.

Goelz LC, David FJ, Sweeney JA, Vaillancourt DE, Poizner H, Metman LV, et al. The effects of unilateral versus bilateral subthalamic nucleus deep brain stimulation on prosaccades and antisaccades in Parkinson’s disease. Exp Brain Res 2017; 235(2): 615–26.

Hoehn MM, Yahr MD. Parkinsonism: onset, progression and mortality. Neurology 1967; 17(5): 427–42.

Hohlefeld FU, Huchzermeyer C, Huebl J, Schneider GH, Brücke C, Schönecker T, et al. Interhemispheric functional interactions between the subthalamic nuclei of patients with Parkinson’s disease. Eur J Neurosci 2014; 40(8): 3273–83.

Hughes AJ, Daniel SE, Kilford L, Lees AJ. Accuracy of clinical diagnosis of idiopathic Parkinson’s disease: a clinico-pathological study of 100 cases. Journal of neurology, neurosurgery, and psychiatry 1992; 55(3): 181–4.

Huss DS, Dallapiazza RF, Shah BB, Harrison MB, Diamond J, Elias WJ. Functional assessment and quality of life in essential tremor with bilateral or unilateral DBS and focused ultrasound thalamotomy. Movement disorders : official journal of the Movement Disorder Society 2015; 30(14): 1937–43.

Knight EJ, Testini P, Min HK, Gibson WS, Gorny KR, Favazza CP, et al. Motor and Nonmotor Circuitry Activation Induced by Subthalamic Nucleus Deep Brain Stimulation in Patients With Parkinson Disease: Intraoperative Functional Magnetic Resonance Imaging for Deep Brain Stimulation. Mayo Clin Proc 2015; 90(6): 773–85.

Kumar R, Lozano AM, Sime E, Halket E, Lang AE. Comparative effects of unilateral and bilateral subthalamic nucleus deep brain stimulation. Neurology 1999; 53(3): 561–6.

Lanciego JL, Luquin N, Obeso JA. Functional neuroanatomy of the basal ganglia. Cold Spring Harb Perspect Med 2012; 2(12): a009621.

Langston JW, Widner H, Goetz CG, Brooks D, Fahn S, Freeman T, et al. Core assessment program for intracerebral transplantations (CAPIT). Movement disorders : official journal of the Movement Disorder Society 1992; 7(1): 2–13.

Le Jeune F, Drapier D, Bourguignon A, Peron J, Mesbah H, Drapier S, et al. Subthalamic nucleus stimulation in Parkinson disease induces apathy: a PET study. Neurology 2009; 73(21): 1746–51.

Lee EM, Kurundkar A, Cutter GR, Huang H, Guthrie BL, Watts RL, et al. Comparison of weight changes following unilateral and staged bilateral STN DBS for advanced PD. Brain Behav 2011; 1(1): 12–8.

Leentjens AF, Dujardin K, Marsh L, Martinez-Martin P, Richard IH, Starkstein SE, et al. Apathy and anhedonia rating scales in Parkinson’s disease: critique and recommendations. Movement disorders : official journal of the Movement Disorder Society 2008; 23(14): 2004–14.

Lhommee E, Klinger H, Thobois S, Schmitt E, Ardouin C, Bichon A, et al. Subthalamic stimulation in Parkinson’s disease: restoring the balance of motivated behaviours. Brain : a journal of neurology 2012; 135(Pt 5): 1463–77.

Lizarraga KJ, Jagid JR, Luca CC. Comparative effects of unilateral and bilateral subthalamic nucleus deep brain stimulation on gait kinematics in Parkinson’s disease: a randomized, blinded study. Journal of neurology 2016; 263(8): 1652–6.

Mansouri A, Taslimi S, Badhiwala JH, Witiw CD, Nassiri F, Odekerken VJJ, et al. Deep brain stimulation for Parkinson’s disease: meta-analysis of results of randomized trials at varying lengths of follow-up. J Neurosurg 2018; 128(4): 1199–213.

Marin RS. Apathy: Concept, Syndrome, Neural Mechanisms, and Treatment. Seminars in clinical neuropsychiatry 1996; 1(4): 304–14.

Marin RS, Biedrzycki RC, Firinciogullari S. Reliability and validity of the Apathy Evaluation Scale. Psychiatry research 1991; 38(2): 143–62.

Mark VW, Oberheu AM, Henderson C, Woods AJ. Ballism after stroke responds to standard physical therapeutic interventions. Arch Phys Med Rehabil 2005; 86(6): 1226–33.

Martinez-Fernandez R, Pelissier P, Quesada JL, Klinger H, Lhommee E, Schmitt E, et al. Postoperative apathy can neutralise benefits in quality of life after subthalamic stimulation for Parkinson’s disease. Journal of neurology, neurosurgery, and psychiatry 2016; 87(3): 311–8.

Montgomery SA, Asberg M. A new depression scale designed to be sensitive to change. The British journal of psychiatry : the journal of mental science 1979; 134: 382–9.

Novak P, Klemp JA, Ridings LW, Lyons KE, Pahwa R, Nazzaro JM. Effect of deep brain stimulation of the subthalamic nucleus upon the contralateral subthalamic nucleus in Parkinson disease. Neurosci Lett 2009; 463(1): 12–6.

Oldfield RC. The assessment and analysis of handedness: the Edinburgh inventory. Neuropsychologia 1971; 9(1): 97–113.

Olsson M, Nikkhah G, Bentlage C, Bjorklund A. Forelimb akinesia in the rat Parkinson model: differential effects of dopamine agonists and nigral transplants as assessed by a new stepping test. The Journal of neuroscience : the official journal of the Society for Neuroscience 1995; 15(5 Pt 2): 3863–75.

Ortega-Cubero S, Clavero P, Irurzun C, Gonzalez-Redondo R, Guridi J, Obeso JA, et al. Effect of deep brain stimulation of the subthalamic nucleus on non-motor fluctuations in Parkinson’s disease: two-years’ follow-up. Parkinsonism & related disorders 2013; 19(5): 543–7.

Paxinos G WC. The rat brain in stereotaxic coordinates. 1998.

Périer C, Agid Y, Hirsch EC, Féger J. Ipsilateral and contralateral subthalamic activity after unilateral dopaminergic lesion. Neuroreport 2000; 11(14): 3275–8.

Rizvi SJ, Pizzagalli DA, Sproule BA, Kennedy SH. Assessing anhedonia in depression: Potentials and pitfalls. Neuroscience and biobehavioral reviews 2016; 65: 21–35.

Robert GH, Le Jeune F, Lozachmeur C, Drapier S, Dondaine T, Peron J, et al. Preoperative factors of apathy in subthalamic stimulated Parkinson disease: a PET study. Neurology 2014; 83(18): 1620–6.

Samii A, Kelly VE, Slimp JC, Shumway-Cook A, Goodkin R. Staged unilateral versus bilateral subthalamic nucleus stimulator implantation in Parkinson disease. Movement disorders : official journal of the Movement Disorder Society 2007; 22(10): 1476–81.

Schwab RS, England AC, Jr. Amantadine HCL (Symmetrel) and its relation to Levo-Dopa in the treatment of Parkinson’s disease. Transactions of the American Neurological Association 1969; 94: 85–90.

Sjoberg RL, Lidman E, Haggstrom B, Hariz MI, Linder J, Fredricks A, et al. Verbal fluency in patients receiving bilateral versus left-sided deep brain stimulation of the subthalamic nucleus for Parkinson’s disease. J Int Neuropsychol Soc 2012; 18(3): 606–11.

Temel Y, Boothman LJ, Blokland A, Magill PJ, Steinbusch HW, Visser-Vandewalle V, et al. Inhibition of 5-HT neuron activity and induction of depressive-like behavior by high-frequency stimulation of the subthalamic nucleus. Proc Natl Acad Sci U S A 2007; 104(43): 17087–92.

Thobois S, Ardouin C, Lhommee E, Klinger H, Lagrange C, Xie J, et al. Non-motor dopamine withdrawal syndrome after surgery for Parkinson’s disease: predictors and underlying mesolimbic denervation. Brain 2010; 133(Pt 4): 1111–27.

Vachez Y, Carcenac C, Magnard R, Kerkerian-Le Goff L, Salin P, Savasta M, et al. Subthalamic Nucleus Stimulation Impairs Motivation: Implication for Apathy in Parkinson’s Disease. Movement disorders : official journal of the Movement Disorder Society 2020; 35(4): 616–28.

Vachez YM, Creed MC. Deep Brain Stimulation of the Subthalamic Nucleus Modulates Reward-Related Behavior: A Systematic Review. Front Hum Neurosci 2020; 14: 578564.

Walker HC, Watts RL, Guthrie S, Wang D, Guthrie BL. Bilateral effects of unilateral subthalamic deep brain stimulation on Parkinson’s disease at 1 year. Neurosurgery 2009; 65(2): 302–9; discussion 9-10.

Walker HC, Watts RL, Schrandt CJ, Huang H, Guthrie SL, Guthrie BL, et al. Activation of subthalamic neurons by contralateral subthalamic deep brain stimulation in Parkinson disease. J Neurophysiol 2011; 105(3): 1112–21.

Welter ML, Houeto JL, Tezenas du Montcel S, Mesnage V, Bonnet AM, Pillon B, et al. Clinical predictive factors of subthalamic stimulation in Parkinson’s disease. Brain : a journal of neurology 2002; 125(Pt 3): 575–83.

Zoon TJ, de Bie RM, Schuurman PR, van den Munckhof P, Denys D, Figee M. Resolution of apathy after dorsal instead of ventral subthalamic deep brain stimulation for Parkinson’s disease. Journal of neurology 2019.

Zoon TJC, van Rooijen G, Balm G, Bergfeld IO, Daams JG, Krack P, et al. Apathy Induced by Subthalamic Nucleus Deep Brain Stimulation in Parkinson’s Disease: A Meta-Analysis. Movement disorders : official journal of the Movement Disorder Society 2020.

Zoon TJC, van Rooijen G, Balm G, Bergfeld IO, Daams JG, Krack P, et al. Apathy Induced by Subthalamic Nucleus Deep Brain Stimulation in Parkinson’s Disease: A Meta-Analysis. Movement disorders : official journal of the Movement Disorder Society 2021; 36(2): 317–26.

